# Hetero-multicellular stromal cells incorporate into scaffold-free 3D cultures of epithelial cancer cells to drive invasion

**DOI:** 10.1101/2025.01.21.634082

**Authors:** Elizabeth Ortiz, Kyaw Hsu Thway, Gabriela Ortiz-Soto, Paulina Yao, Jonathan A Kelber

## Abstract

Breast cancer (BC) is the second leading cause of cancer-related death among women in the U.S. Organoid models of solid tumors have been shown to faithfully recapitulate aspects of cancer progression such as proliferation and invasion. Although patient-derived organoids (PDOs) and patient-derived xenograft organoids (PDXOs) are pathophysiologically relevant, they are costly to propagate, difficult to manipulate and comprised primarily of the most proliferative cell types within the tumor microenvironment (TME). These limitations prevent their use for elucidating cellular mechanisms of disease progression that depend upon tumor-associated stromal cells which are found within the TME and known to contribute to metastasis and therapy resistance. Here, we report on methods for cultivating epithelial-stromal multicellular 3D cultures. Advantages of these methods include a cost-effective system for rapidly generating organoid-like 3D cultures within scaffold-free environments that can be used to track invasion at single-cell resolution within hydrogel scaffolds. Specifically, we demonstrate how to generate these hetero-multicellular 3D cultures using BT-474 breast cancer cells in combination with fibroblasts (BJ-5ta), monocyte-like cells(THP-1) and/or endothelial cells (EA.hy926). Additionally, differential fluorescent labeling of cell populations enables time-lapse microscopy to define 3D culture assembly and invasion dynamics. Notably, the addition of any two stromal cell combinations to 3D cultures of BT-474 cells significantly reduces circularity of the 3D cultures, consistent of the presence of organoid-like or secondary spheroid structures. In tracker dye experiments, fibroblasts and endothelial cells co-localize in the peripheral organoid-like protrusions and are spatially segregated from the primary BT-474 spheroid. Finally, hetero-multicellular 3D cultures of BT-474 cells have increased hydrogel invasion capacity. Since we observed these protrusive structures in hetero-multicellular 3D cultures of both non-tumorigenic and tumorigenic breast epithelial cells, this work provides an efficient and reproducible method for generating organoid-like 3D cultures in a scaffold-free environment for subsequent analyses of phenotypes associated with solid tumor progression.

**SUMMARY:** There is a critical need for 3D cancer models that capture hetero-cellular crosstalk to study cancer metastasis. Our study presents the generation of hetero-multicellular stromal-epithelial in a scaffold and scaffold free environment that can be used to study invasion and cellular spatial distributions.

## INTRODUCTION

Cancer progression is now recognized to be dependent of two major factors: the genetic/epigenetic changes in the tumor cells and a myriad of interaction with non-tumor cells in the tumor microenvironment (TME)^1^. While genetic changes in cells are acknowledged to be necessary for tumor initiation, such alterations alone are not sufficient for tumor progression and metastasis^2^. Components of the TME, originally thought to be silent bystanders, are now known to actively promote cancer progression via mutual and dynamic crosstalk with tumor cells^3^. The TME composition differs depending on the tissue the tumor originates from, the tumor stage, and patient characteristics, but hallmark features include stromal cancer-associated fibroblasts (CAFs), extracellular matrix (ECM), vascular endothelial cells, adaptive and myeloid immune cells^1,4^.

Stromal CAFs in the TME are composed of fibroblast subtypes of diverse origins and functions^5^. Such CAFs are key components of the TME as they interact with tumor cells at several interfaces. CAFs secrete ECM proteins that alters the stiffness of the matrix which may either limit drug delivery via excessive deposition of collagen, proteoglycans and fibronectin or allow tumor cells to invade from the primary tumor site via secretion of ECM degrading matrix metalloproteinases (MMPs)^6,7^. In addition, CAFs promote tumor growth, migration and vascularization via secretion of variety of growth factors, cytokines and angiogenic factors such as epidermal growth factor (EGF), transforming growth factor β(TGF-β) and vascular endothelial growth factor (VEGF) respectively^1,6^. In parallel, endothelial cells, prompted by the hypoxic TME, also promote tumor vascularization and suppress immune cell functions via increased secretion of angiogenic factors and lowered secretion of leukocyte adhesion molecules^1,8^.

With the apparent intricate complexity of cancer progression, it has become essential to incorporate TME stromal components in basic cancer research. However, the establishment of models that faithfully recapitulate known tumor pathophysiology is still a significant unmet need^9,10^. While traditional two-dimensional (2D) cell culture models are easy to handle, rapidly cultured and are highly reproducible, it is comprised of only rapidly proliferating cancer cell clones and do not reflect the cellular heterogeneity found in tumors^10–12^. In a similar manner, transgenic mouse models also do not capture human tumor biology due to low genetic heterogeneity from inbreeding, significant difference in immune system, and histological complexity^13,14^. Due to such limitations, therapeutics developed from classical cancer models often fail to translate to clinical settings.

Patient derived cancer models such as patient derived xenografts and patient derived organoids can address drawbacks of conventional cancer models by capturing in-situ tumor molecular features, genetic background and cellular organization^10,11&15^. However, such patient derived xenografts and organoids require complicated engraftment procedures and long culturing time^16,17^. Combined with variation in tumor acquisition and sampling sites and poor efficiency in cryopreservation, there is a need to develop models that act as a bridge between classical 2D cell cultures and patient derived cancer models^11,18^. In this regard, 3D models of cell culture can serve as models that can be cultured rapidly and capture important in vivo tumor features such as cell-cell interaction, cell-ECM interaction, hypoxia, angiogenesis and production of ECM^19,20^. 3D cell culture models are categorized into scaffold-free and scaffold-based model systems. In scaffold-free systems, cells are induced to self-aggregate into spherical shape by using specific low-attachment cell culture plates or by manipulating the physical parameters of culturing methods. Established methods to obtain scaffold-free 3D spheroids range from simple cell pellet cultures by centrifugation to hanging microplate drops, magnetic levitation and dynamic bioreactor and microfluidic systems^20,21^. Scaffold-based 3D cell cultures are established by addition of polymer-or hydrogel-based scaffolds to imitate the physiological extracellular matrix^19,22^.Such models hold immense potential to model in vivo cellular organization, topology, matrix attachment, migration and drug response.

In addition to scaffold manipulation for extracellular matrix composition models in disease states, 3D cell cultures can also be used to model heterogenous cellular population in a tumor microenvironment. 3D cell cultures composed of cancer cells, and stromal fibroblasts or endothelial cells have been used to study interaction of cancer and individual non-tumor cell lines^23–25^. Reproducible and cost-effective methods for expanding such 3D cell cultures composed of multiple heterogenous cell lines would help researchers elucidate tumor progression. Here, we report on methods for cultivating epithelial-stromal multicellular 3D cultures to study proliferation, invasion, and cell state plasticity. The protocol describes scaffold free and basement membrane solution scaffold-based 3D cultures of breast cancer cells co-cultured with a combination of stromal cells ranging from fibroblasts (BJ-5ta), endothelial (Ea.hy926) cells and monocytes-like cells (THP-1). Breast cancer is currently the second most common cancer worldwide and the most diagnosed cancer in women in the United States^26^. Death from breast cancer is largely due to the metastatic and therapy resistant nature of the disease as overall and metastasis-free survivorship is significantly reduced in patients diagnosed with aggressive HER2 enriched and basal-like breast cancer subtypes^27^. Our described 3D cell culture protocols may assist in development of cost-effective, rapid and reproducible culturing methods that may be paired with formalin-fixed paraffin-embedded tissue preservation methods and subsequent spatial biology applications.

**PROTOCOL:** highlight 3 pages of protocol for filming

1. Cell culture medium

1.1. Cell culture medium for BJ-5ta, BT474, EA.hy926, and MDA-MB-468: Supplement 500mL of Dulbecco’s Modified Eagle Medium (DMEM) high-glucose with 10% heat-inactivated fetal bovine serum (FBS) and 1% penicillin-streptomycin with a pipette. Incorporate 0.1% gentamicin with a micropipette. The medium must be prepared inside a biosafety cabinet.
1.2. Cell culture medium for MCF10A: Supplement 500mL of DMEM/F12 with 5% horse serum and 1% penicillin-streptomycin with a pipette. Incorporate 0.1% gentamicin, 1 mL of 1 µg/mL hydrocortisone, 500 µL of 10 µg/mL insulin, 50 µL of 100 ng/mL cholera toxin, and 10 µL of 20 ng/mL epidermal growth factor with a micropipette. The medium must be prepared inside a biosafety cabinet.
1.3. Cell culture medium for MCF10Ca1h: Supplement 500mL of DMEM/F12 with 5% horse serum and 1% penicillin-streptomycin with a pipette. Incorporate 0.1% gentamicin with a micropipette. The medium must be prepared inside a biosafety cabinet to avoid contamination.
1.4. Cell culture medium for THP-1: Supplement 500mL of RPMI 1640 with 10% FBS and 1% penicillin-streptomycin with a pipette. Incorporate0.1% gentamicin with a micropipette. The medium must be prepared inside a biosafety cabinet to avoid contamination.
1.5. The cell culture medium is changed every 2-3 days for BJ-5ta, BT474, EA.hy926, MDA-MB-468, MCF10A, and MCF10Ca1h until the cells reach 70-80% confluency by assessing their growth daily with a bright field microscope. The cell culture medium is changed once a week for THP-1 cells. The medium must be prepared inside a biosafety cabinet to avoid contamination.
1.6. BJ-5ta, BT474, EA.hy926, MDA-MB-468, MCF10A, and MCF10Ca1h cells are grown in 100 mm Dish Cell Culture Treated Surface in standard cell culture incubators with 5% CO2.
1.7. THP-1 cells are grown T75 Cell Culture Treated Flasks in standard cell culture incubators with 5% CO2.
2. Cell collection

2.1. Turn on the UV light to sanitize the interior of the biosafety cabinet for 15 min.
2.2. Open the biosafety cabinet window sash to stabilize the airflow and turn on the vacuum aspiration system.
2.3. Clean the interior hood surface and the tubing of the vacuum aspiration system with 70% ethanol.
2.4. Prepare fresh serum-free cell culture medium inside the biosafety cabinet: supplement 500mL of Dulbecco’s modified Eagle’s medium (DMEM) high-glucose with 10% heat-inactivated fetal bovine serum (FBS) and 1% penicillin/streptomycin with a pipette. Incorporate 0.1% Gentamicin with a micropipette.
2.5. Warm the cell culture medium to 37°C, phosphate-buffered saline (PBS), and Trypsin-EDTA (0.25%) by placing the items in a bead bath before starting the experiment.
2.6. Ensure that the cells are 70-80% confluent through visual inspection through a microscope.
2.7. Aspirate and discard the culture medium from the plated cells with the vacuum aspirator. Wash the remaining medium once with 2 mL of PBS with a pipette then aspirate and discard the PBS with the vacuum aspirator.
NOTE: THP-1 cells are grown in suspension. Steps 7-9 can be omitted.
2.8. Add 1 mL of Trypsin in the cell plate with a micropipette and place the plate inside a 5% CO2incubator at 37°C for 5 min.
2.9. Inactivate the trypsin by adding 1 mL of defined trypsin inhibitor to the plate with a micropipette. Disperse cell clusters by pipetting the liquid mixture with a P1000 micropipette and collect the cell solution from the bottom of the plate.
Transfer the cell suspension with a micropipette to a 15 mL conical tube and centrifuge at 100 x g for 5 min at room temperature. Discard the supernatant with the vacuum aspirator.
2.10. Continue to cell counting.
3. Preparation of working dye solution and staining cells in suspension

3.1. Before opening the vial of cell tracker molecular fluorescent probes (see table of materials), allow the product to warm to room temperature for 15 min in a bead bath set at 37°C.
3.2. Dissolve the lyophilized cell tracker blue dye (mass = 5mg, molecular weight = 209.6 g/mol) to a final concentration of 10mM with 2.385 mL of DMSO using a micropipette.
3.3. Dissolve the lyophilized cell tracker orange dye (mass = 50ug, molecular weight = 550.4 g/mol) to a final concentration of 10mM with 9.084 µL of DMSO using a micropipette.
3.4. Dissolve the lyophilized cell tracker deep red dye (mass = 15 ug, molecular weight = 698.3 g/mol) to a final concentration of 1 mM with 20 µL of DMSO using a micropipette.
3.5. Prepare the working cell tracker blue dye media solution (5 µM) by diluting 1 µL of the dye in 2 mL of serum-free DMEM medium with a micropipette.
3.6. Prepare the working cell tracker orange dye media solution (5 µM) by diluting 1 µL of the dye in 2 mL of serum-free DMEM medium with a micropipette.
3.7. Prepare the working cell tracker deep red dye media solution (1 µM) by diluting 2 µL of the dye in 2 mL of serum-free DMEM medium with a micropipette.
3.8. Resuspend the epithelial cells, BJ-5ta fibroblasts and Ea.hy926 endothelial cells in the prepared working cell tracker blue, orange, and deep red dye media solution (2mL) respectively with a micropipette.
3.9. Incubate the tubes at 37°C in the 5% CO_2_ incubator for 30 min.
3.10. After 30 min of incubation, centrifuge the tubes at 100x g for 5 min at room temperature.
3.11. Aspirate and discard the supernatant with the vacuum aspirator and resuspend the pellet thoroughly in 1mL of 10% FBS serum containing DMEM medium with a micropipette.
3.12. Continue to cell counting.
4. Cell Counting

4.1. Collect 10 µL of the cell suspension and transfer it to a microtube with a micropipette.
4.2. Mix with 10 µL of Trypan blue and pipette thoroughly.
4.3. Transfer 20µL of the cell-trypan solution with a micropipette to a cell counting chamber slide. Insert and count the cells using an automated cell counter.
4.4. Calculate the average total live cell number from two readings.
5. Calculations

5.1. Working cell stocks must be prepared for each cell type at a concentration of 6.67x10^3 cells/mL equivalent to 2,000 cells/300µL. The total stock volume will depend on the experimental sample size.
For mono-culture spheroids, each independent sample will consist of 300 µL of the epithelial cell working stock equivalent to 2,000 epithelial cells.
5.2. For co-culture spheroids (two cell types), each independent sample will consist of 150 µL of epithelial cell working stock and 150 µL of stromal cell working stock. The sample will contain 1,000 epithelial cells and 1,000 stromal cells.
5.3. For co-culture spheroids (three cell types), each independent sample will consist of 150 µL of epithelial cell working stock, 75 µL of stromal cell #1 working stock, and 75 µL of stromal cell #2 working stock. The sample will contain 1,000 epithelial cells, 500 stromal cells of working stock #1, and 500 stromal cells of working stock #2.
6. Plating

6.1. Transfer the required volume with a micropiette for 3 technical replicates, plus 1 extra, into a microtube. Mix thoroughly with a pipette, then transfer 300 µL of the sample to a well of a U-shaped-bottom, ultra-low attachment 96-well microplate.
6.2. Repeat the process for each additional technical replicate.
6.3. Place the 96-well plate in an incubator at 37°C.
7. Brightfield Imaging

7.1. Observe spheroid growth and morphology every 24 h up to 96 h using a microscope.
7.2. xImage spheroids using a phase-contrast microscope.
8. Widefield imaging protocol setup for spheroids stained with tracker dyes and spheroids overlayed with basement membrane solution.

8.1. Using the imaging and automated incubator devices listed in the table of materials, turn on the devices and create a new imaging protocol in the task manager of the imaging software. Under the "Procedure" tab, set the automated incubator temperature set point to 37°C and select "Preheat before continuing with the next step."
NOTE: An alternative fluorescent imager can be used if it has the appropriate filters for each of the dyes (blue, deep red and orange) used.
8.2. Set the image settings to the following specifications: Magnification: 4X PL FL Phase, Field of View: 3185 X 3185 µm, Full WFOV
8.3. The specifications for the channels are as follows: DAPI: 377/447 nm, illumination = 10, integration time = 107 ms, gain = 10, RFP: 531/593 nm, illumination = 10, integration time = 137 ms, gain = 10, CY5: 628/685 nm, illumination = 10, integration time = 137 ms, gain = 10
8.4. For imaging spheroids in a basement membrane solution overlay, use the Brightfield specifications: illumination = 10, integration time = 5 ms, and gain = 17.1.
8.5. Select the desired wells for imaging and approve the specification changes under the select wells icon.
8.6. Navigate to the "Data Reduction" tab to adjust the cellular analysis settings.
8.7. Set the threshold value to 19,500 with a light background and select "Fill holes in masks."
8.8. For object selection, set the minimum object size to 100 µm and the maximum object size to 1000 µm, and select "Analyze entire image."
8.9. Only a primary mask and object count are needed for this analysis.
8.10. Save all changes and open the imager application.
8.11. Place the experimental U-shaped-bottom microplate into the incubator by opening the drawer, then close it using the imager software.
8.12. To run the protocol, select the “Procedure Info” tab, add a user, and choose the protocol.
8.13. Ensure the correct plate type is selected and set the imaging time to 30 min per plate.
8.14. Select the desired imaging interval, indicate whether the plate has a lid, and adjust the imaging start time and duration.
8.15. Click "Schedule Plate/Vessel" to start the imaging process.
9. Basement membrane solution overlay (optional)

9.1. Basement membrane solution can be applied to spheroids 24 h post-plating.
9.2. Fill an ice bucket with ice to keep the basement membrane solution cold and store it at 4°C when not in use.
9.3. Aspirate approximately 170 µL of medium with a multichannel pipette.
9.4. Use a magnifying glass and a mini lightbox to closely observe the small spheroids. Place the 96-well spheroid plate over the lightbox and position the magnifying glass overhead.
9.5. Set the P200 pipette or alternative to 30µL and collect basement membrane extract to create three microdroplets.
9.6. Add one microdroplet over each spheroid of the three technical replicates with a micropipette.
9.7. Ensure the 96-well plate is flat and position the pipette vertically above the spheroid. Release the droplet without touching the bottom of the well.
9.8. Place the plate in an incubator at 37°C for 20 min.
9.9. Overlay the spheroids with an additional 50 µL of basement membrane solution per well with a micropipette.
9.10. Reposition spheroids to the center of the well using a pipette tip.
9.11. Incubate at 37°C for 30 min.
9.12. Add 100 µL of cell culture medium to each well.
10. Quantification

10.1. Download and open the ImageJ software. Upload the spheroid images to ImageJ.
10.2. Set Measurements: Go to Analyze > Set Measurements and select "Area" and "Centroid." Measure Spheroids: Use the lasso tool to trace the spheroid and measure the object.
10.3. Collect Data: Record the values for three biological replicates and calculate the average area and circularity.
10.4. Convert the area from pixels to square meters using the following formula: Area (m²) = ((550/504) × √ (area in pixels)) ²
10.5. Analyze Data Using GraphPad Prism:

10.5.1. Select "Enter replicate values, stacked into columns."
10.5.2. Input the data into the columns and select the monoculture spheroid control and the co-culture treatment spheroid samples.
10.5.3. Perform a one-way ANOVA column analysis.
10.6. ANOVA Parameters:

10.6.1. No matching or pairing.
10.6.2. Assume a Gaussian distribution of residuals.
10.6.3. Assume equal standard deviations.
10.6.4. Use Tukey’s multiple comparison test to compare the mean of each column with the mean of every other column.
11. Widefield Immunofluorescent Image Processing

11.1. Save the images directly from the imager software as PNG or Tiff file format.
11.2. Select each individual channel or overlap channels and save as PNG file.
11.3. Adjust brightness and contrast in the imager software or in ImageJ if images are exported as Tiff files

## REPRESENTATIVE RESULTS

In this study, we developed a cell culture system to generate hetero-multicellular 3D spheroids consisting of epithelial and stromal cells with organoid-like morphology. Spheroids were established via plating 2000 epithelial cells in monoculture conditions. In co-culture conditions of two cell types, spheroids were established via plating 1000 epithelial cells and 1000 stromal cells. In coculture conditions of three cell types, spheroids were established via plating 1000 epithelial cells and 2 different stromal cell types of 500 cells each cell type. Upstream of spheroid establishment, cells can be stained with fluorescent cell tracker dyes that allows for monitoring cellular spatial organization. After 24 h of initial spheroid formations, downstream applications include pharmacological perturbations, imaging, and sample collection. Time-lapse imaging is useful for assessing changes in spheroid behavior and morphology, including area and circularity (Figure 1a). At 24 h post-plating, spheroids can be embedded in a scaffold environment, and time-lapse imaging can be used to assess the onset of invasive structures from the spheroid (Figure 1b). The collection of hetero-multicellular spheroid samples has many applications, including genomic and proteomic profiling at the global and single-cell levels through experimental techniques such as RNA sequencing, single-cell RNA sequencing, proteomic sequencing, and cyclic immunofluorescence.

**Figure 1:**
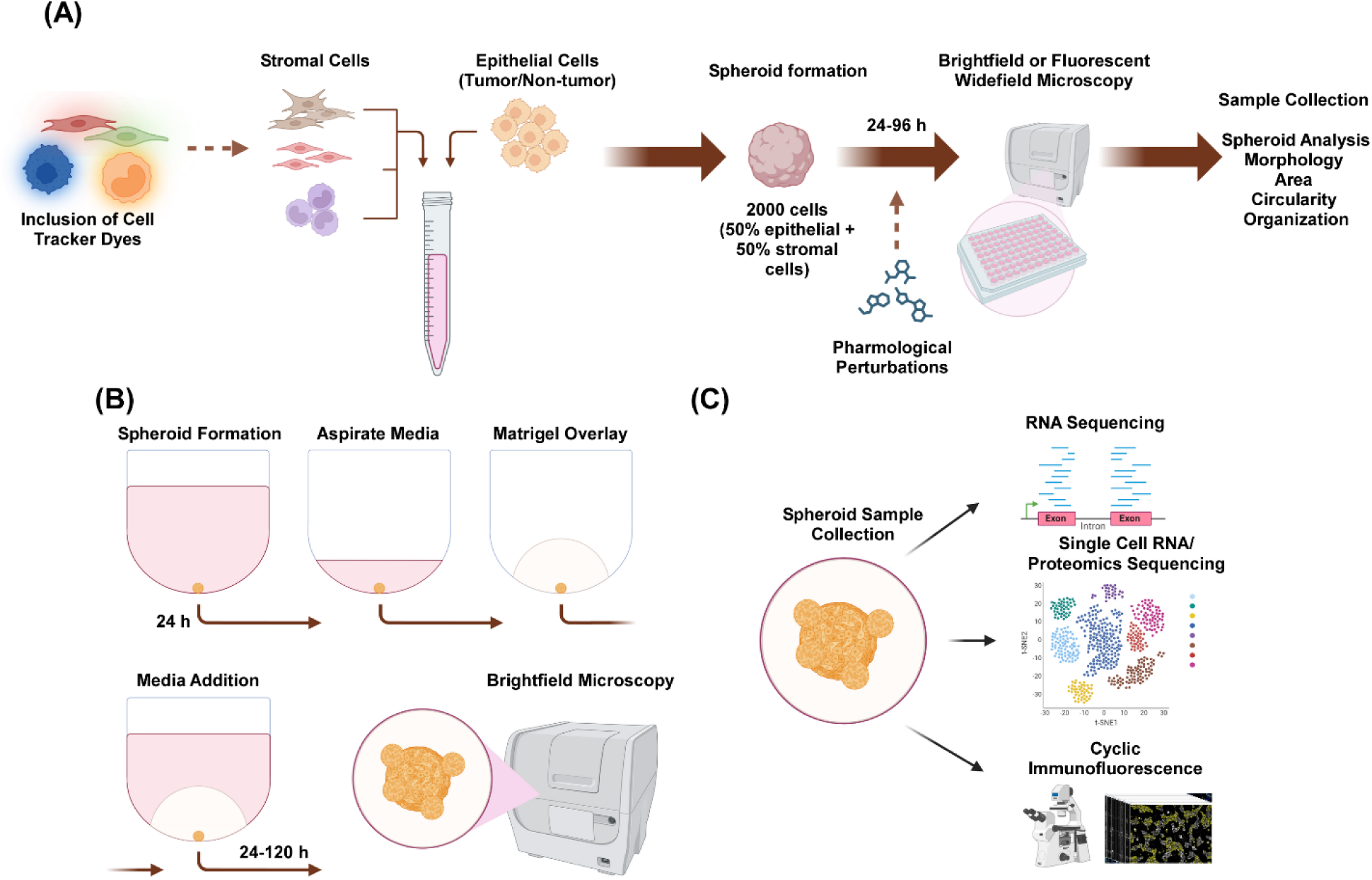
Schematic representation of the 3D cell culture process and potential applications. (a) Cell suspension of epithelial cells with non-epithelial stromal cells is pipetted into 3D ultra-low attachment plates to form spheroids. Spheroids are imaged in Brightfield microscopy every 24 h across 96 h. Cell lines can be stained with fluorescent cell tracker dyes for downstream widefield microscopy before inducing spheroid formation or can be perturbed pharmacologically after spheroid formation. Spheroid parameters such as morphology, area, circularity, and organization can be analyzed. (b) Spheroids of epithelial and epithelial cells with stromal cells are formed using the protocol from (a). Spheroids were established via plating 2000 epithelial cells in monoculture conditions. In co-culture conditions of two cell types, spheroids were established via plating 1000 epithelial cells and 1000 stromal cells. In coculture conditions of three cell types, spheroids were established via plating 1000 epithelial cells and 2 different stromal cell types of 500 cells each cell type. After 24 h, a scaffold like basement membrane solution is overlaid, and images are captured in brightfield microscopy every 24 h for 120 h. (c) Established scaffold-free and scaffold-based hetero-multicellular 3D cultures can be used for a variety of downstream applications such as cyclic immunofluorescence, single-cell RNA sequencing and single-cell proteomics.

MCF10A, MCF10Ca1h, and BT-474 monoculture spheroids maintain a compact spherical phenotype for up to 96 h post-plating. When co-cultured with EA.hy926 microvascular endothelial cells, BJ-5ta fibroblasts, and/or THP-1 monocyte-like cells, the spheroids developing cellular protrusions at the periphery, which becomes more pronounced at 96 h (Figure 2a, b, c). The budding morphology ranges from solid to loose cell aggregates, resembling organoid morphology. In contrast, MDA-MB-468 monoculture spheroids appear as large, loose cell aggregates. However, when MDA-MB-468 cells are co-cultured with EA.hy926, BJ-5ta, and/or THP-1, they form compact spheroids (Figure 2d).

**Figure 2:**
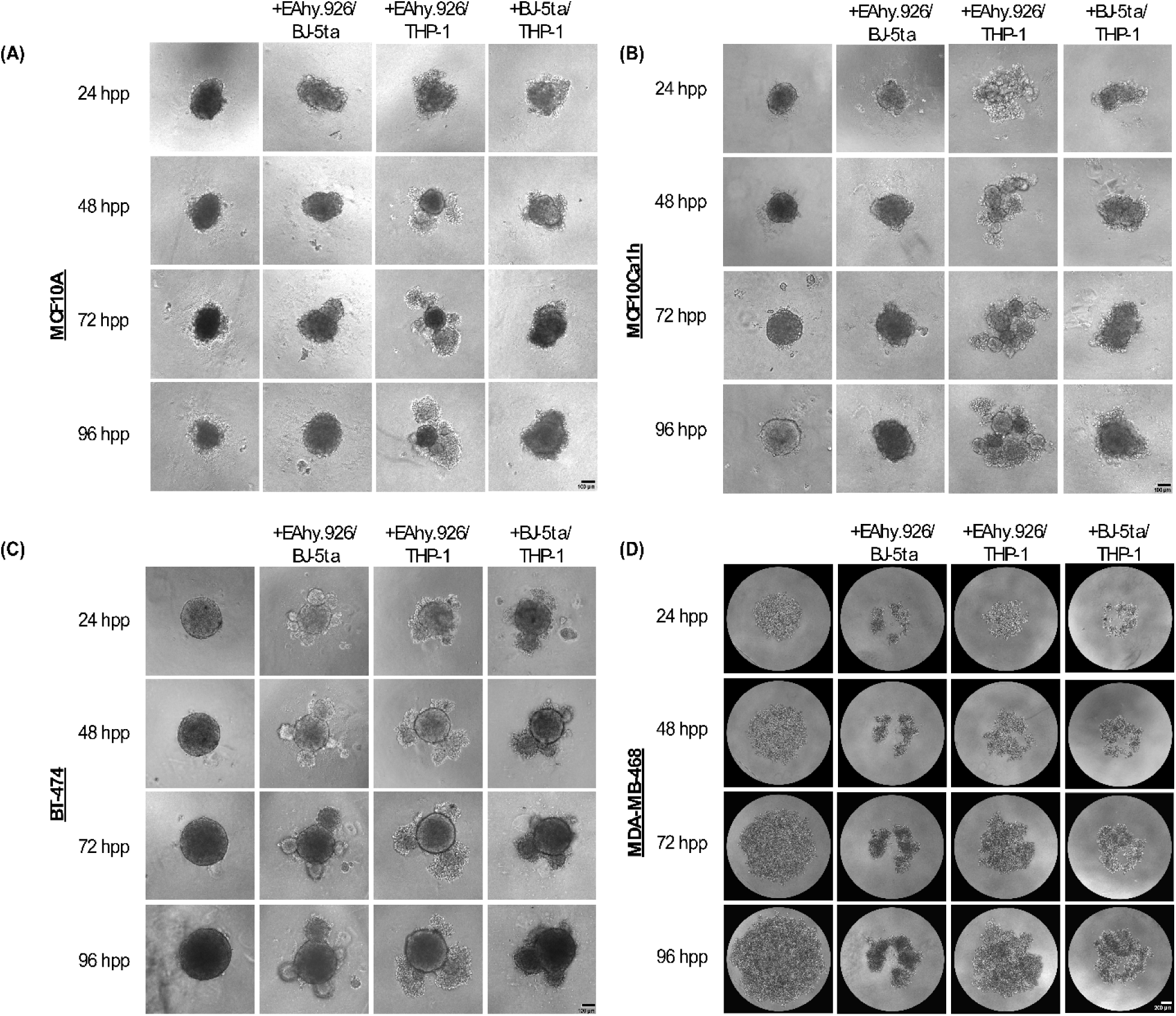
Non-epithelial stromal cells adopt a range of architectures & morphologies when combined with epithelial tumor/non tumor cells in scaffold-free 3D cultures. Representative brightfield images of (a) MCF10A (b) MCF10Ca1h (c) BT-474 (d) MDA-MB-468 spheroids in monoculture or in co-culture conditions with stromal BJ-5ta fibroblasts/Ea.hy926 microvascular endothelial cells or Ea.hy926/THP-1 monocyte-like cell or BJ-5ta/THP-1 cells across 96 h. Each spheroid was formed by plating 2,000 cells. Spheroids in monoculture conditions were formed using 2,000 epithelial cells. Spheroids in co-culture conditions were formed using 1,000 epithelial cells and 2 different stromal cell types of 500 cells. Formation of budding or aggregated organoid-like structures is initiated in spheroid co-culture conditions 24 h post plating in MCF10A, MCF10Ca1h and BT-474 epithelial cells. Compaction of MDA-MB-468 cells is observed in spheroid co-culture conditions 25 h post plating. hpp = hours post plating

At 72 h post-plating, MCF10A, MCF10Ca1h, and BT-474 cells co-cultured with EA.hy926 and THP-1, or with BJ-5ta and THP-1, exhibited a significant increase in spheroid area compared to monoculture epithelial spheroids. MCF10Ca1h also showed a significant increase in spheroid area when co-cultured with EA.hy926 and BJ-5ta. The onset of budding structures in co-cultured spheroids led to a significant decrease in spheroid circularity for MCF10A, MCF10Ca1h, and BT-474 co-cultured with EA.hy926 and THP-1, or with BJ-5ta and THP-1. Similar effects were observed for BT-474 co-cultured with both EA.hy926 and BJ-5ta (Figure 3a, b). In contrast, MDA-MB-468 cells co-cultured with EA.hy926 and BJ-5ta, EA.hy926 and THP-1, or BJ-5ta and THP-1, showed a significant decrease in spheroid area compared to monoculture MDA-MB-468 spheroids yet no effect was observed in circularity (Figure 3a, b).

**Figure 3:**
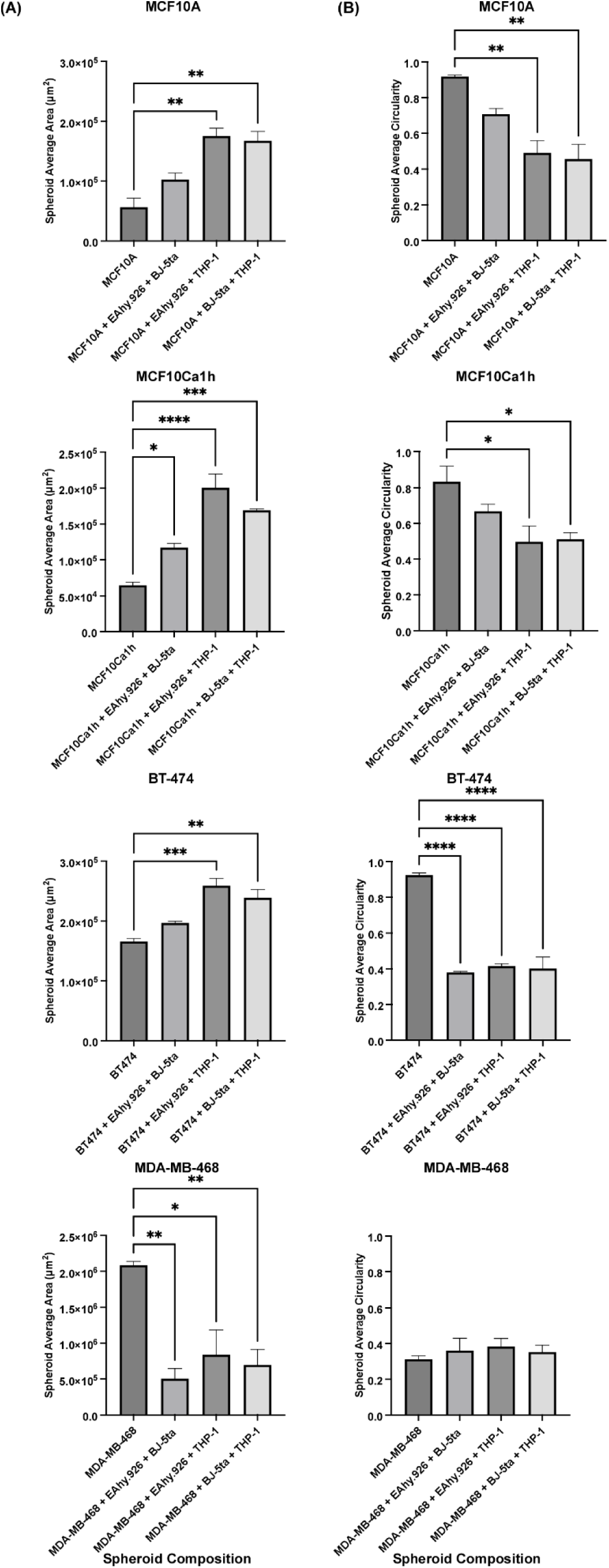
Area and circularity analysis graphs of mono-cultured and hetero-multicellular stromal 3D scaffold-free cultures at 72 h post-plating. (a) Average area (cm2) and (b) Average Circularity of MCF10A, MCF10Ca1h, BT-474, and MDA-MB-468 in mono-cultured and hetero-multicellular stromal spheroid cultures 72 h post plating. Data reported are representative of at least 3 independent biological replicates and are reported as technical replicate averages ± SEM, unless otherwise indicated. *, **, *** or **** represent *p* values < 0.05, 0.01, 0.001, or 0.0001 respectively, unless otherwise noted.

Application of cell tracker dye to BT-474 tumorigenic epithelial cells and stromal cells prior to spheroid establishment demonstrated that stromal cells, including EA.hy926 and BJ-5ta form the budding structures at the perimeter of the central BT474 spheroids (Figure 4, Supplementary Videos 1-4). At 48 h post plating, individual widefield fluorescent images in spheroids co-cultured with fibroblasts reveal fibroblast spheroids co-localize together with endothelial cells but do not co-localize with BT-474 spheroids. A minority of endothelial cells are also found to co-localize with BT-474 spheroids in co-culture conditions. This suggests that the arrangement of stromal cells within the spheroid is correlated with an organoid-like morphology.

**Figure 4:**
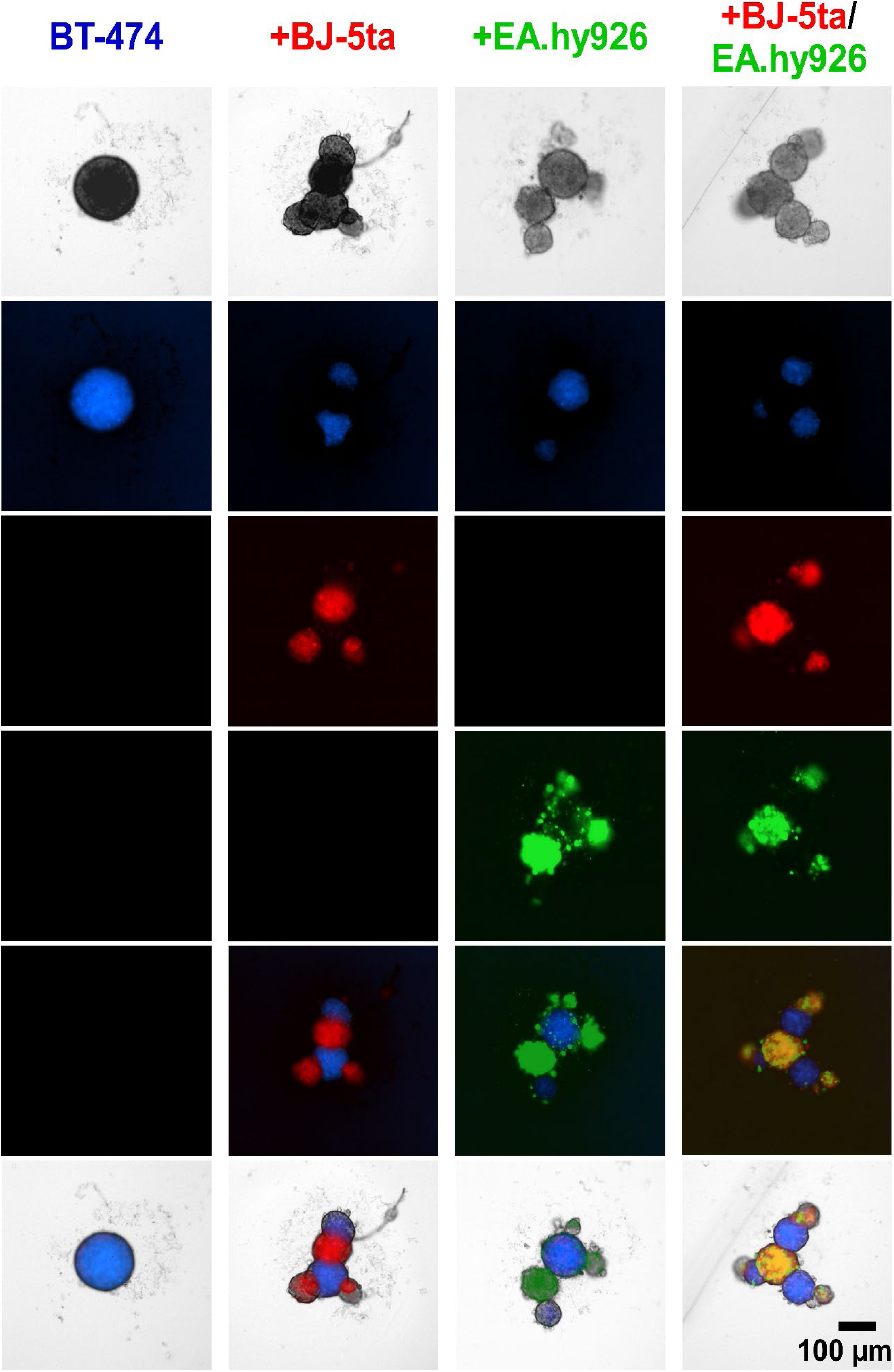
Wide-field fluorescence still images of differentially dyed stromal and BT-474 cells in hetero-multicellular 3D cultures at 48 h post-plating. BT-474 spheroids are stained with blue cell tracker fluorescent dye. BJ-5ta fibroblasts are stained with orange cell tracker dye, represented in red color. Ea.hy926 endothelial cells are stained with deep red cell tracker dye, represented in green color. Each spheroid was formed by plating 2,000 cells. Spheroids in monoculture conditions were formed using 2,000 epithelial cells. Spheroids in co-culture conditions (BT-474/BJ-5ta & BT-474/Ea.hy926) were formed using 1,000 epithelial cells and 1,000 stromal cells. Spheroids in double co-culture conditions (BT-474/BJ-5ta/Ea.hy926) were formed using 1,000 epithelial cells and 500 cells of each stromal cell type. Figures are representatives of at least 3 biological replicates.

To assess the biological relevance of our organoid spheroid model, spheroids were overlaid with basement membrane solution 24 h after plating. Monoculture BT-474 spheroids displayed no invasive properties 120 h post-plating. However, BT-474 spheroids co-cultured with BJ-5ta or EA.hy926 developed structures at the periphery of the spheroid, which invaded the scaffold basement membrane solution environment. The number and length of these protrusions were significantly enhanced in BT-474 spheroids co-cultured with both BJ-5ta and EA.hy926 by 48 h through 120 h post-plating. (Figure 5, Supplementary Videos 5-8)

**Figure 5:**
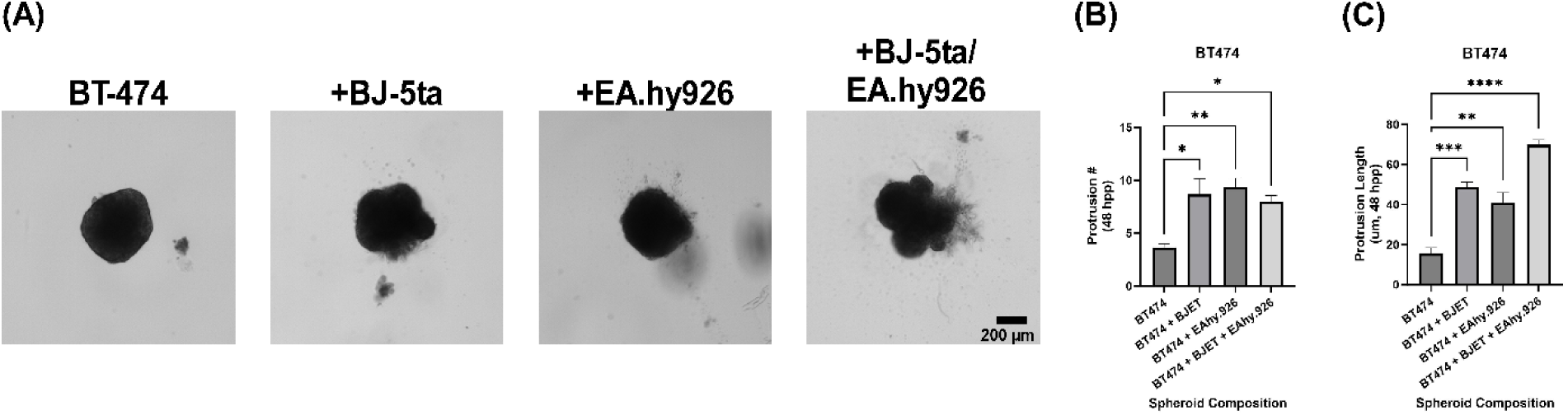
Brightfield still images of BT-474 cells in hetero-multicellular 3D cultures with basement membrane solution overlay at 5 days post-overlay. Each spheroid was formed by plating 2,000 cells. basement membrane solution is overlaid over spheroids 24 h post plating. Spheroids in monoculture conditions were formed using 2,000 epithelial cells. Spheroids in co-culture conditions (BT-474/BJ-5ta & BT-474/Ea.hy926) were formed using 1,000 epithelial cells and 1,000 stromal cells. Spheroids in double co-culture conditions (BT-474/BJ-5ta/Ea.hy926) were formed using 1,000 epithelial cells and 500 cells of each stromal cell type. A) Invasive structures protruding from the cancer spheroid embedded in basement membrane solution can be observed in co-culture conditions. B) Qualification of invasive protrusion count at time 48 hpp. C) Qualification of invasive protrusion length at time 48 hpp.Figures are representatives of at least 3 biological replicates. hpp = hours post plating

## Supplementary Video Captions

**Supplementary Video 1.1:** BT-474 monoculture spheroid with cell tracker blue dye. 2,000 BT-474 cells were resuspended in a Nuclon Sphera ULA plates to form spheroids. An overlap of brightfield and widefield fluorescent images are captured across 48 h.

**Supplementary Video 1.2:** BT-474 spheroids co-cultured with BJ-5ta fibroblasts. 1,000 BT-474 and 1,000 BJ-5ta fibroblasts were resuspended in a Nuclon Sphera ULA plates to form spheroids. BT-474 cells were incubated with cell tracker blue dye and BJ-5ta fibroblasts were incubated with cell tracker orange dye. An overlap of brightfield and widefield fluorescent (blue = BT-474, red = BJ-5ta) images are captured across 48 h.

**Supplementary Video 1.3:** BT-474 spheroids co-cultured with Ea.hy926 endothelial cells. 1,000 BT-474 and 1,000 Ea.hy926 endothelial cells were resuspended in a Nuclon Sphera ULA plates to form spheroids. BT-474 cells were incubated with cell tracker blue dye and Ea.hy926 endothelial cells were incubated with cell tracker deep red dye. An overlap of brightfield and widefield fluorescent (blue = BT-474, green = Ea.hy926) images are captured across 48 h.

**Supplementary Video 1.4:** BT-474 spheroids co-cultured with BJ-5ta fibroblasts and Ea.hy926 endothelial cells. 1,000 BT-474, 500 BJ-5ta fibroblasts and 500 Ea.hy926 endothelial cells were resuspended in a Nuclon Sphera ULA plates to form spheroids. BT-474 cells were incubated with cell tracker blue dye. BJ-5ta fibroblasts and Ea.hy926 endothelial cells were incubated with cell tracker orange dye and deep red dye respectively. An overlap of brightfield and widefield fluorescent (blue = BT-474, red = BJ-5ta, green = Ea.hy926) images are captured across 48 h.

**Supplementary Video 2.1:** BT-474 monoculture spheroid in basement membrane solution. 2,000 BT-474 cells were resuspended in a Nuclon Sphera ULA plates to form spheroids and the spheroids were embedded in a basement membrane solution 24 h post plating. Brightfield images were taken across 60 h.

**Supplementary Video 2.2:** BT-474 spheroids co-cultured with BJ-5ta fibroblasts in basement membrane solution. 1,000 BT-474 and 1,000 BJ-5ta fibroblasts were resuspended in a Nuclon Sphera ULA plates to form spheroids and the spheroids were emebbed in a basement membrane solution 24 h post plating. Brightfield images were taken across 60 h.

**Supplementary Video 2.3:** BT-474 spheroids co-cultured with Ea.hy926 endothelial cells in basement membrane solution. 1,000 BT-474 and 1,000 Ea.hy926 endothelial cells were resuspended in a Nuclon Sphera ULA plates to form spheroids and the spheroids were embedded in a basement membrane solution 24 h post plating. Brightfield images were taken across 60 h.

**Supplementary Video 2.4:** BT-474 spheroids co-cultured with BJ-5ta fibroblasts and Ea.hy926 endothelial cells in basement membrane solution. 1,000 BT-474, 500 BJ-5ta fibroblasts and 500 Ea.hy926 endothelial cells were resuspended in a Nuclon Sphera ULA plates to form spheroids and they spheroids were embedded in a basement membrane solution 24 h post plating. Brightfield images were taken across 60 h.

## DISCUSSION

Our hetero-multicellular spheroid model demonstrates that epithelial-stromal cell interactions drive stromal cell budding in scaffold-free 3D culture conditions and the formation of invasive structures in scaffold-based 3D culture conditions. We observed consistent budding structure formation in both tumorigenic cell lines (MCF10Ca1h and BT-474) and the non-tumorigenic epithelial cell line MCF10A (Fig. 2a-c). Interestingly, while spheroids from cell lines MCF10A, MCF10Ca1h, and BT-474, whether in monoculture or co-culture with fibroblasts and endothelial cells, formed circular and compact budding structures in scaffold-free environments, spheroids from cell line MDA-MB-468 showed less compaction (Fig. 2d). This was unexpected, as MDA-MB-468 cells were anticipated to form circular spheroids, like the other tested epithelial cell lines. Cell line SUM-149, a hybrid epithelial-mesenchymal primary breast cancer cell line, was found to form compact spheroids like MCF10A, MCF10Ca1h, and BT-474 (data not shown)^28^. Future research should investigate the signaling pathways mediating the formation of less compact spheroids in the MDA-MB-468 cell line.

Regarding the combination of different cell lines, THP-1 monocyte-like cellsinduce formation of loose aggregate structures in all co-culture conditions of spheroids. Less compaction is observed in budding structures in co-culture conditions with THP-1/Ea.hy926 cells compared to those in co-culture conditions with THP-1/BJ-5ta cells (Fig 2a-d). Therefore, the compaction of spheroids is influenced by the stromal cell composition in the model^29,30^.

Inconsistent plating of the hetero-cell combinations can result in incomplete spheroid development or variability in spheroid size, as compared to their technical replicates. Ensuring consistency in cell plating is crucial, as the spatial arrangement of cells that determines spheroid phenotype and development occurs primarily within the first 48 h^31^. During this critical period, cell ratios play a significant role, with the proportion of stromal cells affecting both spheroid area and circularity.

Loose spheroid aggregates, which are dependent on cell type, can easily disassemble during manual handling. However, manual manipulation is necessary when overlaying spheroids with a scaffold-like basement membrane solution or when collecting whole spheroid samples. To mitigate this issue, it is advisable to work at a slower pace when collecting spheroids via pipette aspiration and to minimize manual handling whenever possible. During the basement membrane solution overlay, spheroids are prone to shifting within the well, often settling at the well’s edge. A basement membrane solution-to-media ratio is important for maintaining the matrix properties of the 3D system; however, the spheroids cannot be fully enveloped due to the viscosity of the basement membrane solution, so manual intervention is required^32^. Non centered spheroid positioning poses a challenge for imaging, as light refraction near the plastic edge of the well can interfere with image clarity.

The spheroids used in this study were composed of human cancer cell lines, and the methods have not yet been applied to primary cell lines. Our results suggest that specific cell-cell interactions can drive the formation of organoid-like structures, indicating the potential applicability of this method to a wide range of tumorigenic cell lines. However, while our model attempts to mimic the complexity of the tumor microenvironment, the differing rates of cellular proliferation present a challenge for long-term studies^33^. Rapidly proliferating cells tend to outgrow slower proliferating ones, making this model less suitable for extended experiments. In addition, different growth media type requirements for cell lines can introduce additional factors that may lead to spheroid phenotypes. Most cell lines in the study were grown in DMEM media. Studies with cell lines grown in different media may require different culture conditions testing a variety of media ratios, yet this effect is observed in long term experimentation^30^.

Our model offers several advantages over other 3D cancer models. The spheroids we generate have a uniform and reproducible initial morphology and size, making it easier to compare treatment outcomes with controls. In contrast, some models using single cells suspended in media or scaffolds can result in variable cellular spatial distributions, leading to inconsistency. Our model demonstrates that interactions between distinct cell populations can be effectively studied.

Traditionally, organoid and spheroid models have been composed of monoculture cancer cells or cancer cell co-cultures to investigate gradients of nutrients, hypoxia, or cellular arrangement^34^. Additionally, our model can be studied in a scaffolded environment within 24 h post-plating, providing an opportunity to explore early invasion events driven by diverse cell-cell interactions. While alternative 3D cancer approaches employ different scaffold matrix components to study changes in spheroid morphology and invasiveness, our model complements these by allowing investigation of these changes in the context of both cell population dynamics and scaffold composition^35^.

Most spheroids produced using our methods retain their structural integrity after manual handling, allowing for sample collection in downstream applications. Conventional genetic modifications such as CRISPR-Cas9, lentiviral shRNA transduction and siRNA interference can be introduced upstream of spheroid formation. The complex behaviors observed in these spheroids suggest dynamic alterations in gene expression, which can be further investigated using RNA-seq. Advanced techniques such as CycIF, scRNA-seq, CosMx, and Visium can also be employed to study genomics and proteomics at spatial or single-cell levels.

This spheroid model has the potential to mimic *in vivo* studies of targeted tumor therapeutics by capturing interactions between stromal cell buds and the therapy before it reaches the cancer cells. This is significant because it can help determine whether stromal cells influence the drug’s potency, potentially protecting the cancer cells from the therapeutic effects.

## Supporting information

Supp Vid 1.1

Supp Vid 1.2

Supp Vid 1.3

Supp Vid 1.4

Supp Vid 2.1

Supp Vid 2.2

Supp Vid 2.3

Supp Vid 2.4

## ACKNOWLEDGMENTS

We thank members of the Baylor University Developmental Oncogene Laboratory for their helpful remarks and feedback during preparation of this manuscript. Funding support was provided by Baylor University Department of Biology and College of Arts and Sciences, NIH-NIGMS 2SC1GM121182 (to J.A.K.).

